# Speed Synchrony Promotes Collective Motion in Mixed-Species Fish Schools

**DOI:** 10.64898/2026.07.10.737720

**Authors:** Jahanvi Tiwari, Arshed Nabeel, Viraj Torsekar, Juee Dhar, Fathimath Lamshana, Vishwesha Guttal

## Abstract

Principles of collective motion are now well established, though research has largely focused on homogeneous groups. Heterogeneity is widespread in animal groups, e.g. arising from sex, size or even species, raising a central question: can collective behaviour emerge when individuals have distinct behaviours? Here, we combine experiments and modelling to investigate mixed-species collective motion using two closely related fish species, rosy barbs and tiger barbs. In conspecific groups, both species exhibit collective motion, but they differ strikingly in their intrinsic movement: tiger barbs exhibit slow- and fast-swimming, whereas rosy barbs display fast swimming only. Despite this difference, these species readily form mixed-species schools where the slow swimming speed of tiger barbs disappears, and the collective motion is dominated by a single fast-swimming mode. We develop an individual-based model incorporating local interactions involving speed matching. Our model demonstrates that bimodal speed in conspecific schools of tiger barbs is an emergent property that is lost in mixed-species groups. Additionally, despite high cohesion, we observe spatial sorting of the two species within the mixed-species groups, which our model explains through differences in inter- and intra-specific interactions. Our results provide experimental evidence that canonical principles of collective motion extend to heterogeneous mixed-species groups.

## Introduction

In animal groups, heterogeneity arises from differences in size, variations in behavioural tendencies, or social roles such as leadership, sex, or even species identity [1, 2, 3, 4]. Additionally, biologists have documented a wide variety of so-called mixed-species groups, in which members of more than one species form a group and move together [5, 6, 7, 8]. This heterogeneity can potentially alter the nature of local interactions: for example, individuals may interact differentially with neighbours based on leadership, dominance, kinship, or species identity, leading to asymmetric or non-reciprocal interaction strengths [9, 10]. However, much of the studies in the field of collective motion, including experiments and models, have largely focused on homogeneous groups, treating all individuals as identical with uniform interaction rules [11, 12, 13, 14, 15, 16, 17, 18, 19]. Incorporating heterogeneity into experimental and theoretical frameworks of collective behaviour is not only biologically realistic, but also potentially important to physicists and engineers working on swarm dynamics [20, 1, 2, 21].

Mixed-species groups are compelling natural systems for investigating the role of heterogeneity in collective behaviour. Mixed-species flocks are found in a wide variety, ranging from weak aggregates to highly coordinated mobile flocks [5]. However, much of the literature on mixed-species groups focuses on functions and adaptations via field-based studies [5, 7], with only a few investigating the structure and dynamics of heterogeneous flocks from a quantitative perspective [22, 23, 24, 25, 26]. A central question in this context is to understand how heterogeneity in behavioural interactions affects the tendency of two or more species to group and move together. To maintain group coherence, individuals must coordinate their activity and adjust their behaviour in response to the movement behaviours of other group members [13, 14, 27], a process broadly referred to as activity matching [28, 5]. Activity matching can be further quantified as speed matching, in which individuals try to match their speed to the group’s. Most flocking models assume speed is constant (but see [29, 30]) and thus miss out on speed matching, an important principle underlying coordination among individuals. Using the lens of collective motion, we expect that simple local interactions among individuals, such as a tendency to match the speed and direction of motion with nearby conspecifics and heterospecifics, can facilitate heterospecific schooling. Whether this expectation holds true even when the participating individuals differ in their intrinsic movement dynamics, as is likely the case with many heterospecific groups, remains unclear. Testing these ideas requires a precise quantitative characterisation of collective motion dynamics.

Here, we investigate the structural and dynamical properties of free-swimming mixed-species schools of two closely related barb species under laboratory conditions. The controlled laboratory conditions enable us to use high-resolution tracking and quantitative characterization of the collective motion of mixed-species groups, which is otherwise daunting under field conditions. The two species we study exhibit dramatically different speed profiles within conspecific groups but show highly coordinated movement in heterospecific groups, providing a unique system for studying the emergent properties of mixed-species groups. We develop an individual-based model that incorporates speed variability and elucidates how simple local interactions, such as speed matching, can facilitate heterospecific schooling and reproduce key features of the experimental data.

## Results

We study mixed-species schooling behaviour in juveniles of two closely related fish species, tiger barbs (*Puntius tetrazona*) and rosy barbs (*Pethia conchonius*). Both fish species readily show collective motion under laboratory conditions (Fig. 1) and are known compatible tank-mates. Rosy barbs are native to South Asia, whereas tiger barbs are native to Southeast Asia. However, due to the wide prevalence of tiger barbs in the aquarium trade and some reported cases of introduction to the wild, these barb species occur geographically overlapping in many parts of the world [31, 32], with tiger barbs being a potential invasive species.

**Figure 1.**
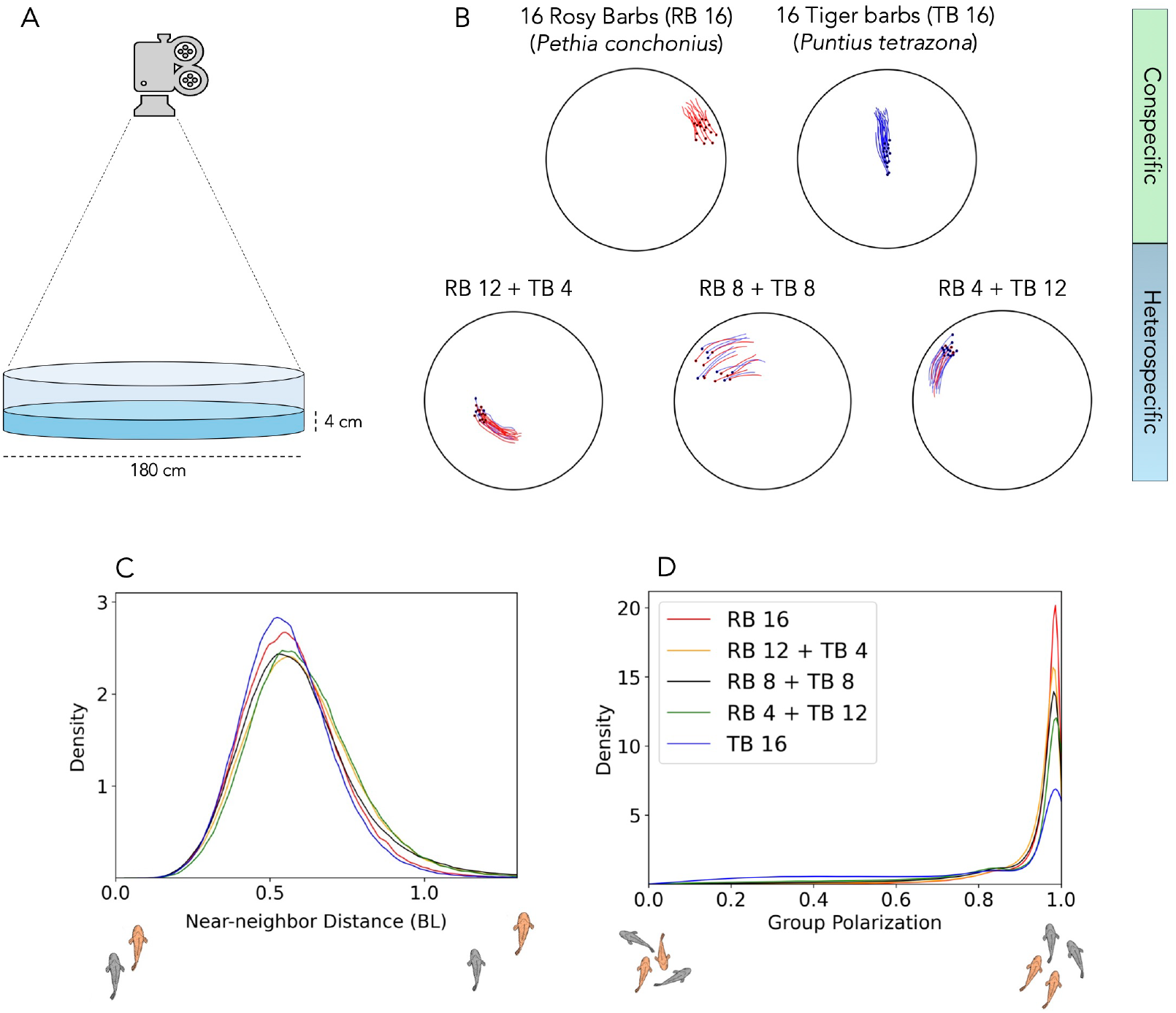
Experimental set-up for mixed species schooling behaviour. (A) Using an overhead high-resolution camera, we record fish schools in a circular arena of 180 cm diameter with shallow water of 4 cm depth. (B) Experimental conditions involved conspecific schools of 16 rosy barbs (RB 16), 16 tiger barbs (TB 16), and three treatments of heterospecific schools of 16 fish (the bottom row); dots represent sample instantaneous positions of fish and lines represent sample trajectories, with red and blue colours representing rosy barbs and tiger barbs, respectively. In all treatments, fish readily form cohesive groups and exhibit collective motion. (C) shows the distributions of near-neighbour distances, a measure of group cohesion. (D) shows the distribution group polarisation, a measure of group alignment.

Our experimental treatments correspond to five group compositions, of which two were conspecific groups of each species and three were mixed species groups of the two species with varying ratios of each (Fig. 1 B). Consistent with many field studies showing that fish schools are naturally size sorted [33, 34, 35, 36, 37], the body size variation was kept minimal across experiments, with a mean body length of 3.7 cm and an average coefficient of variation of 6.1% (*SI Appendix*, Table. S1). We convert all units from cm to mean body length (BL) to normalize measurements across trials. In our experiments, we recorded the free-swimming behaviour of groups of 16 fish in a circular arena of 180 cm diameter under shallow water conditions (4 cm)(Fig. 1 A), a strategy typically used in many experimental studies to confine fish movement to effectively two dimensions [38, 15]. Each experiment was recorded at 25 frames per second lasting 90 minutes, and each treatment was replicated 5 times (one replicate of TB 16 was discarded due to unreliable tracking). Fish were not reused across different experiments. We extracted fish trajectories from recorded videos using TRex [39]. See Materials and Methods section for detailed experimental protocols.

### High degree of cohesion and alignment in mixed species schools

We observe that both species readily show collective motion in conspecific as well as mixed species group compositions (Fig. 1 B-D and *SI Appendix*). We characterize the degree of group cohesion via the near-neighbour distance (NND) computed for each fish. We find an average splitting percentage of less than 10% per hour across all treatments, indicating that even heterospecific schools are highly cohesive (*SI Appendix*, Fig. S1). Since group splitting is rare, NND is a good measure of cohesion, where smaller NND indicates high cohesion and larger NND value indicates low cohesion. We find that the groups exhibit a high degree of cohesion, with NND distributions showing a mode at around 0.5 body lengths regardless of the species composition of the group (Fig. 1 C); see *SI Appendix*, Table. S2 for additional summary statistics of the distributions. Further, the low degree of splitting and high cohesion also makes it evident that there is no differential space use between the species: all individuals mostly swim together as one cohesive group throughout the duration of every trial.

We quantified group alignment via polarisation 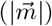, defined as the magnitude of the average orientation vector across all fish. We find that the groups were highly aligned, with a mode of group polarisation close to the maximum value of 1, for both conspecific and heterospecific schools (Fig. 1D).

### Speed matching facilitates heterospecific schooling

A key necessary condition for the formation and maintenance of heterospecific grouping is that group members match their activity with each other [5, 28]. In the context of collective movement, we expect that fish match their speed to that of their group members [23]. To test this, we compute and compare individual speed distributions across treatments. First, we find a striking difference between the types of movements in conspecific schools of rosy barbs and tiger barbs. Rosy barbs exhibit a unimodal distribution of individual speed (mode at 8.1 BL/s, Fig. 2A) corresponding to a fast swimming behavior. On the other hand, tiger barbs show a bimodal speed distribution (one mode at 1.1 BL/s and the other at 7.3 BL/s, Fig. 2B), indicating a slow and a fast swimming behaviour. Importantly, we find that bimodality is not a consequence of trial to trial variation in fish behaviour; further, we also confirm that this is not due to within trial behavioural variation over time (*SI Appendix*, Fig. S2). Therefore, rosy barbs and tiger barbs provides an interesting system to study heterospecific schooling, since the conspecific groups of these species show contrasting speed profiles.

**Figure 2.**
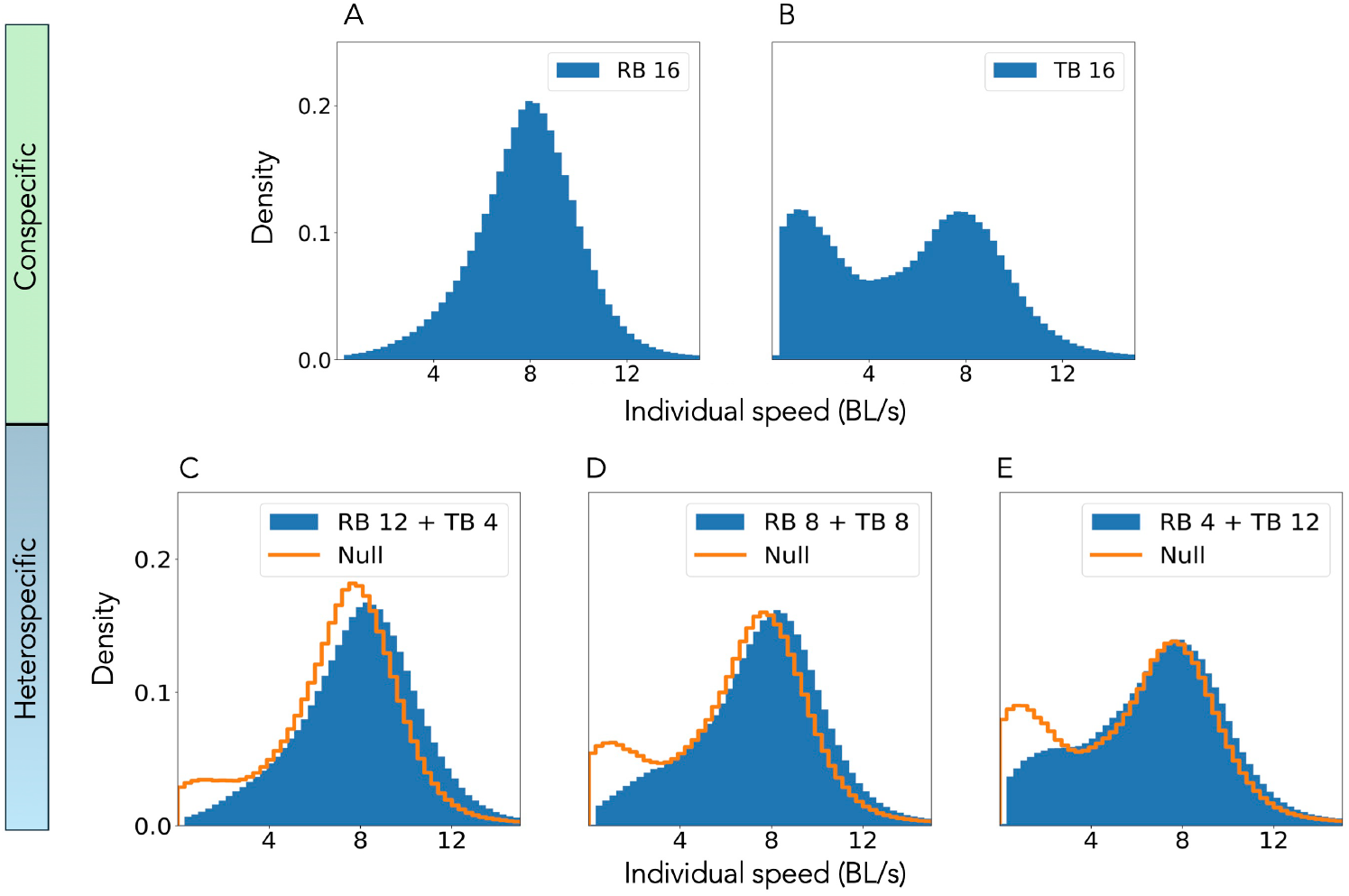
Speed synchrony in mixed species schools. Rosy barbs show a unimodal distribution of individual speeds (A), while tiger barbs show a bimodal distribution of speeds (B). In all heterospecific treatments (C to E), fish show unimodal distributions of speeds. The null distributions (dotted lines) are constructed from the weighted ratios of speed distributions from conspecific schools. The bimodality expected from the null data is absent in the experimental heterospecific treatments, providing evidence for speed matching and movement synchrony between the two species.

In the mixed-species groups, we mainly observe the fast-swimming behaviour across all treatments (Fig. 2 C-E). We plot the null data for each heterospecific group obtained by pooling the speed data from the two conspecific groups, weighted by their respective ratio in the mixed-species group. Based on the null data, we expect a bimodal distribution for individual speed in at least two of the three heterospecific group treatments (Fig. 2. D-E). Clearly, we do not observe bimodality in the speed distributions for mixed-species treatments. Furthermore, the observed mode of the speed distribution of fish in heterospecific groups is in the range 7.9 to 8.3 BL/s, which remarkably coincides with the fast swimming mode of rosy barb conspecific groups at ≈ 8.1 BL/s (*SI Appendix*, Table. S2). In other words, one species (tiger barbs) *appears* to lose their slow-swimming behaviour, resulting in matching their activity with that of the other species (rosy barbs). These results provide evidence for the activity matching hypothesis, i.e. individuals in mixed species groups match their activity (speed) to maintain group cohesion. Is this apparent activity matching a consequence of tiger barbs actively modifying their behaviour depending on the group composition? Or, is there a parsimonious explanation for the apparent activity matching, that does not include explicit modification of intrinsic behavior by tiger barbs? In the upcoming sections, we will explore this question using an individual-based model.

### Spatial sorting of species within the mixed species group

In heterogeneous schools, assortment based on size, speed, familiarity [34, 12] and even species [24, 40, 41] is often reported. Hence, we ask whether individuals of same species assort into conspecific subgroups while still maintaining group cohesion and alignment with heterospecifics. To quantify this, we calculate strong sorting percent which measures how frequently conspecifics form a highly connected part of the heterospecific group. We then randomize the sorting data 1,000 times to create a null distribution which can be compared to the observed sorting in experiments. We find that observed strong sorting percent in all the experiments does not overlap with the null distribution (Fig. 3B). In fact, it is significantly higher than the null distribution (*SI Appendix*, Table. S3). Our analysis show that fish do form subgroups of conspecifics within heterospecific groups while maintaining high degree of cohesion and alignment!

**Figure 3.**
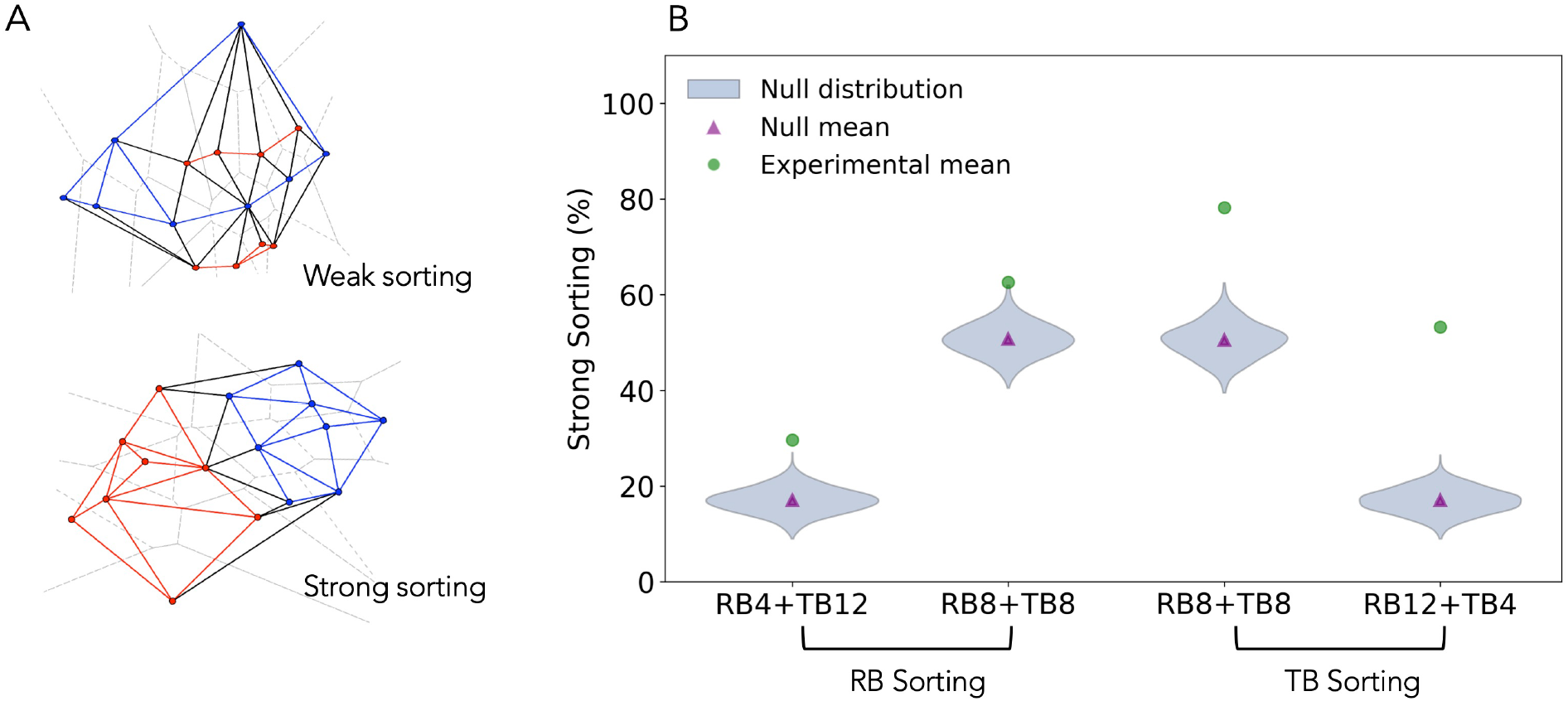
Fish sort based on species identity within heterospecific schools. (A) Voronoi neighbourhood graph of sample experimental snapshots, showing weak and strong sorting; red and blue dots represent rosy barbs and tiger barbs, respectively. If all individuals of the same species of a heterospecific school are connected via Voronoi neighbourhood, we consider this strong sorting. (B) Strong sorting per cent of Rosy barbs and Tiger barbs across different heterospecific treatments. The null distribution was generated by randomly shuffling individual identities within each frame (1,000 permutations). For all the treatments, *p <* 0.001 based on Z-scores, indicating a significantly higher sorting in experimental data (*SI Appendix*, Table. S3).

### A simple individual-based model explains observed patterns in mixed species schools

To better understand the observed patterns of speed bimodality in conspecific groups, speed matching and spatial sorting in mixed-species schools, we build an individual-based, spatially explicit, variable-speed model of collective movement. In our model, an important consideration is that individuals exhibit variable speeds [30, 29, 42]. In addition to interacting with each other through directional *alignment* and *attraction/repulsion* interactions, we consider a new mechanism of interaction via explicit speed-matching: Specifically, fish regulate their speed to match the local group velocity of their neighbourhood, with an interaction rate *µ*_gr_, while also having a preferred swimming speed v_0_ to which they relax with rate *µ*_ind_.

In the mixed-species version of the model, the interaction strengths can have different values for the two species, making the interactions *non-reciprocal* [43]. A detailed description of the model and parameters is given in the *Materials and Methods* section. Our model can produce cohesive and polarised schools, a baseline feature observed in both conspecific and heterospecific groups of both species (see Table. 1 for parameter values, *SI Appendix*, Fig. S4).

**Table 1.** Model parameters used for the two species

| Parameter | Symbol | Rosy barbs | Tiger barbs |
| --- | --- | --- | --- |
| Preferred speed (BL/s) | $v_0$ | 8 | 8 |
| Repulsion zone (BL) | $r_0$ | 0.5 | 0.5 |
| Strength of local alignment (BL/s<sup>2</sup>) | $\mu_{\text{al}}$ | 5 | 5 |
| Strength of speed relaxation (1/s) | $\mu_{\text{ind}}$ | 1 | 1 |
| Strength of local speed matching (1/s) | $\mu_{\text{gr}}$ | 1 | 6 |
| Intraspecific attraction (1/s<sup>2</sup>) | $\mu_{\text{at}}^{\text{intra}}$ | 2 | 6 |
| Interspecific attraction (1/s<sup>2</sup>) | $\mu_{\text{at}}^{\text{inter}}$ | 1 | 1 |
| Noise strength – speed (BL<sup>2</sup>/s<sup>3</sup>) | $D_v$ | 12 | 12 |
| Noise strength – angle (BL<sup>2</sup>/s) | $D_\varphi$ | 12 | 12 |

Our first goal is to explain the nontrivial speed distribution patterns observed in empirical data (Fig. 2). Here, we emphasize that we do not encode the observed bimodality of speeds at the individual level. Rather, our goal is to find the conditions under which such a bimodality in speed emerges. We find that the distribution of individual speeds in conspecific groups is determined by the balance between *µ*_gr_ (the strength of the speed matching interaction introduced in this model) and *µ*_ind_ (rate of relaxation to preferred individual speed). When *µ*_gr_ is larger than *µ*_ind_, i.e. the tendency to match the neighborhood speed is higher than the desire to swim at the preferred speed, the speed distribution becomes bimodal, analogous to the bimodal speed distribution of conspecific groups of tiger barbs (*SI Appendix*, Fig. S5). Conversely, when *µ*_ind_ and *µ*_gr_ are comparable, the speed distribution will be unimodal, analogous to the group behavior of rosy barbs (Fig. 4A). We emphasise that speed bimodality is an emergent property of the groups and is not an in-built feature of the model at the individual level.

**Figure 4.**
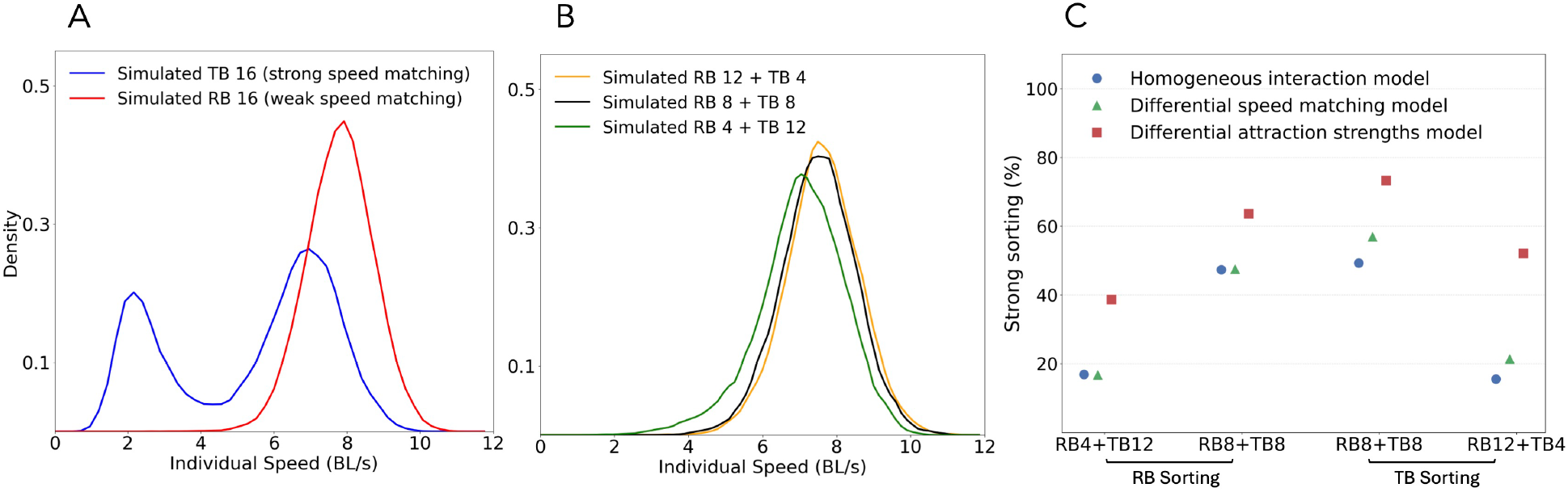
A variable speed individual-based model captures key results of experiments. (A) Single species simulations: Strong speed matching (high *µ*_*g*_ ) in model reproduces the speed bimodality observed in tiger barb schools (blue lines) whereas weak speed matching (low *µ*_*g*_ ) results in unimodal speed distributions observed in rosy barb schools. (B) Mixed-species simulations: In mixed-species simulations, the slow-swimming mode in speed distributions is lost, matching heterospecific treatments from experiments (C) Species sorting within mixed species simulations: In the homogeneous interaction model (blue dots), all interaction strengths are equal across the two species, representing a null scenario of sorting across treatments. The differential speed-matching model – with different *µ*_*g*_ for different species – alone cannot explain the observed spatial sorting (compare green triangles with blue circles). In the differential attraction strengths model (red squares), we include, in addition to differential speed matching, differences in the strengths of intra- and interspecific attractions. This model best captures the spatial sorting (red squares) as observed in the data (Fig. 3 (B)). For parameter values refer to Table 1 in Methods section.

When these ‘virtual’ tiger barbs and rosy barbs are mixed in different proportions, the speed distribution profiles in the simulated mixed-species groups also exhibits a loss of bimodality (Fig. 4B), a key result from our experiments (Fig. 2C-E). Importantly, we are able to reproduce these results without any changes in the parameter values of interaction strengths and with no additional assumptions–such as an inter-species leadership hierarchy– about either species in mixed-species groups. In summary, the model demonstrate that tiger-barbs do not need to adjust their intrinsic properties such as interaction strengths with group composition. Speed matching emerges as a consequence of rosy barbs’ general tendency to swim faster as a group, and tiger barbs’ propensity to match speed to their neighbourhood.

Our second goal is to explain the spatial sorting observed in our experiments (Fig. 3). Previous computational modeling studies demonstrate that differential movement or differential interaction parameters affect the spatial positioning of individuals within mobile groups [12, 44, 10, 9]. So far, we have demonstrated speed matching between two fish species, ruling out differential speed as a possible mechanism for the observed sorting. We now consider the possibility of differential local attraction interactions leading to spatial sorting. Specifically, to explain spatial sorting, we hypothesise that intra-specific attraction interactions to be stronger than inter-specific attraction interactions. Thus, we incorporate separate values for the strength of attraction for intra-specific and inter-specific interactions. Our model simulations demonstrate that making intra-specific attraction stronger than inter-specific attraction is sufficient to reproduce the spatial sorting patterns observed in experimental data (Fig. 4C, *SI Appendix*, Fig. S6). Further, we find that the difference in speed matching rates alone is not sufficient to capture the observed spatial sorting patterns (green triangles in Fig. 4C). We also note that introducing these differential inter- and intra-specific interactions does not alter the patterns of speed profiles.

Put together, we conclude that (i) tiger barbs exhibit a stronger tendency to match local group speed than rosy barbs, and (ii) both species exhibit stronger intra-specific attraction in comparison to inter-specific attraction (see Table. 1). With these assumptions, our model reproduces the key features of both conspecific and heterospecific schooling experiments.

## Discussion

We investigate heterospecific schooling in a controlled experimental setting, allowing us to quantitatively characterize mixed-species groups and reveal their collective properties. We find that despite the close relatedness, tiger barbs and rosy barbs exhibit markedly contrasting speed profiles in conspecific schools. A central result of our study is experimental evidence for speed matching between species, leading to highly cohesive and aligned heterospecific schooling despite these intrinsic differences in speed. Secondly, while being highly cohesive and aligned, heterospecific schools exhibit spatial sorting, i.e., a preferential positioning of fish near conspecifics. We develop a variable-speed agent-based model of heterospecific groups that not only reproduces various features of the data but also reveals a broader principle: local activity (speed and orientation) matching between individuals is a crucial proximate mechanism that facilitates the formation and maintenance of mixed-species schools. Importantly, our model reveals that speed-matching can arise in mixed species schools, without individuals of either species needing to explicitly modify their intrinsic behavioral rules based on their schooling partners.

In conspecific schools, rosy barbs show a unimodal distribution of speed, whereas tiger barbs exhibit a bimodal speed distribution (Fig. 2A-B), suggesting the importance of incorporating variable speed in models of collective motion [29, 30, 42, 45, 46, 8]. To explain these results, we develop a variable-speed model, in which we assume that individuals not only exhibit local directional alignment but also try to match the local group speed. When the propensity of fish to match the group speed is comparable to their relaxation rate to their preferred speed, the model predicts a unimodal speed distribution. On the other hand, an additional mode – corresponding to a slower speed – emerges spontaneously when the fish tend to match the speed of the group much more strongly than their own preferred speed. Thus, our model suggests that tiger barbs have a stronger tendency to follow the group’s speed, whereas rosy barbs seem to balance their preferred individual speed with the group’s speed equally (Fig. 4A). Although some studies have reported a correlation between speed and polarisation in fish schools [15, 42, 27, 47, 21], a feature consistent with our data (*SI Appendix*, Fig. S3), the behaviour of tiger barbs is novel due to its emergent bimodal speed distribution. We emphasise that our model produces highly cohesive and aligned schools under both parameter conditions, consistent with the real data (*SI Appendix*, Fig. S4).

Our experiments show that heterospecific groups of rosy barbs and tiger barbs exhibit high cohesion and alignment (Fig. 1C-D), with a unimodal speed distribution (Fig. 2C-E). A notable observation is that even when rosy barbs are present at a relatively smaller proportion (Fig. 2E), the speed distribution in the mixed species groups becomes unimodal, which is a property of conspecific rosy barb schools. Could this indicate that rosy barbs are dominant partners in heterospecific schooling interactions? Our model provides evidence to the contrary. The model shows that the experimental result of the loss of speed bimodality in heterospecific schools is a natural outcome of local interactions among fish. It is important to note that in our model, we did not assume that tiger barbs preferentially follow rosy barbs. Thus, there is no in-built dominance hierarchy among group members in the model that causes the loss of bimodality. Although similarity in species speeds in heterospecific schools has been recognised in the past [48, 23], our study highlights a dramatic example of the loss of slow swimming speed as an emergent property of heterospecific groups.

Our experiments also show evidence for spatial sorting of the two species based on species identity (Fig. 3B). Spatial sorting, where individuals form sub-groups with conspecifics, has been hypothesized in mixed-species groups, to mitigate costs associated with mixed species grouping, such as cost of coordination and oddity effects [28, 40, 49]. Our model shows that when intra-specific social interactions are stronger than inter-specific interactions (see Table. 1), different species will spontaneously spatially sort within the mixed species school (Fig. 4C), thus offering a potential local interaction mechanism that explains the observed spatial positioning of fish.

In nature, mixed-species groups occur along a spectrum ranging from weakly aggregated mixed-species groups to highly cohesive and aligned well-mixed groups; in the latter extreme, the species identity within a group may not matter [5, 28]. Seen in this context, our experiments show that tiger barbs and rosy barbs form mixed-species schools that exhibit high cohesion and alignment, yet also show preferential spatial positioning based on species identity within the moving groups. While group properties like cohesion and group alignment provide potential anti-predator benefits, sorting within groups can be an additional strategy to counter the oddity effect [50, 49, 24]. We conjecture that the phylogenetic and phenotypic similarities between tiger barbs and rosy barbs may explain why they readily form mixed-species groups. Despite the differences in their native geographic distributions, the two species have some geographically overlapping introduced populations [32, 31]. Furthermore, tiger barb is categorized as a moderately invasive species in some of these introduced populations [31]. Based on our experiments, we predict that even introduced species could potentially form mixed species schools with phenotypically similar species in their introduced habitats. Future studies can investigate whether such an ability to form mixed species groups may provide adaptive benefits in introduced habitats, thereby increasing establishment success and invasive potential.

Broadly, our work demonstrates that the canonical principles and models of collective movement extend to mixed species schools as well. Our computational model was able to reproduce many key experimental results while incorporating fairly minimal assumptions on fish movement and interactions, thus providing a theoretical framework for investigating the structure and dynamics of mixed-species schools. More broadly, we expect that phenotypic similarity and phylogenetic relatedness can play an important role in determining whether two species will form a cohesive and aligned group [5, 51]. As the phenotypic distance between species increases, their ability to school together can be compromised by coordination costs [52]. Future studies may consider how phenotypic and phylogenetic distances influence heterospecific schooling dynamics using experimental setups and integrate these findings with both modeling and field studies of mixed-species groups. We reiterate that much of the collective motion research is conducted in the context of conspecific groups, while the literature on mixed-species groups typically ignores collective motion. We hope that our study inspires more work on integrating the fields of collective motion and mixed-species flocking systems.

## Materials and Methods

### Experimental Set Up

Both the species of fish, tiger barbs (*Puntius tetrazona*) and rosy barbs (*Pethia conchonius*), were obtained from local aquarium vendors, where they are often kept together as tank-mates. The two species are closely related belonging to subfamily Smiliogastrinae and are found in freshwater benthopelagic habitat, feeding on worms, crustaceans, insects and plant matter [53, 54]. Both species have been widely used in aquarium trades throughout the world, with overlapping introduced populations observed in the wild, e.g. in parts of Sri Lanka, Puerto Rico, Singapore and the Magdalena River in Colombia [31, 32].

In laboratory, fish were maintained at 27±1^°^C temperature and sufficient aeration for at least one week before the starting of experiments, which allows them to habituate in the new environmental conditions. The fish were fed every morning at 10 am, except on the day of experiment, where they are fed only after the experiments. We explicitly use fish of similar sizes in our experiments with an average coefficient of variation (CV) in body length across all experiments being less than 6% (*SI Appendix*, Table. S1).

We recorded free-swimming behavior of fish inside a white circular arena. The circular geometry of the arena provides symmetric space with no corners. The arena was filled to a depth of 4 cm water level which is sufficiently shallow water to constrain the motion of the fish to effectively two dimensions and minimise occlusion events. The arena was covered with white curtains on all sides to avoid any external stimulus or perturbations. A video camera (Sony PXW FS5) was mounted at a height of 180 cm above the arena to record the movement of freely swimming fish; the videos were recorded at the spatial resolution of 1920×1080 (35 Mbps) and a temporal resolution of 0.04s (25 frames per second). The radius of the arena was 90 cm to provide sufficient space for movement while facilitating high resolution tracking and minimizing boundary effects. The lighting was homogenized by placing four LED panels symmetrically around the arena (Fig. 1).

To investigate the schooling behavior with varying species composition, we considered conspecific groups of both species and heterospecific groups with varying species composition (25, 50 and 75 percent of each species) for group size 16. Five replicates were done for all the group compositions (one replicate of the treatment TB 16 was not useful due to unreliable tracking), and the order of treatment for replicate was randomized. The number of replicates per treatment is comparable to typical collective behavior studies [29, 15, 55].

Fish were moved from their holding tank to an “acclimation tank” where they spend about 20-24 hours with their group members. The acclimation tanks were maintained at 27 ± 1^°^ C temperature and with sufficient aeration. The fish group was then transferred to the experimental arena, where the water is maintained at the same temperature as the acclimation tank. Temperature, pH and TDS of the water were recorded before and after the trial. The curtains were closed around the setup, and video recording was initiated. For each trial, videos were recorded for approximately 90 minutes. The fish were then transferred to a post experiment holding tank, and the water in the experimental arena was replaced for the next trial to avoid any potential chemical cues from previous trial. All the fish were used only once for the experiments.

### Data Analysis

#### Automated tracking and data extraction

High-resolution fish trajectories were extracted from the videos using automated tracking software TRex [39]. The trajectories were obtained as time-series of the positions of each individual fish, throughout the experimental duration. Although individual fish identities are not maintained throughout the tracks, this is not important for our analysis based on aggregate group parameters typically computed across one frame.

#### Data cleaning and preprocessing

Individual fish trajectories were cleaned by filtering out detections outside the tank boundaries (signifying detection artifacts), and detections showing abnormally high velocities (of 20 BL/s and more, signifying identity-swaps across fish which skews instantaneous speed estimates). We also removed individuals which split from the main group. To identify splitting, we construct a neighbourhood graph for the fish school, with 6 times the median nearest-neighbour distance (NND) as the threshold distance for two fish being connected. We retain only largest connected component in the neighbourhood graph and remove the remaining fish. The exact value of the distance threshold does not have a qualitative impact on the qualitative nature of our results. Finally, we only retain frames where at least 8 fish remain as valid detections after the cleaning procedure, discarding the rest. We save the cleaned trajectories (individual positions over time) after preprocessing, and compute the individual velocities as **v**_*i*_(t) = (**x**_*i*_(t + Δt) ™ **x**_*i*_(t))/Δt, where Δ = 0.04 s. We convert the units from cm to body lengths (BL) to ensure consistency across experiments.

#### Speed, polarisation and nearest-neighbour distance

The magnitude of the individual velocity vector **v**_*i*_(t) gives the individual speed v_*i*_(t), and we define the individual orientation as **e**_*i*_ = **v**_*i*_/v_*i*_.

The group polarization vector **m** is computed as

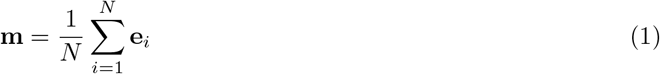

The magnitude of the polarisation vector, |**m**|, determines the degree of order in the school, with 0 denoting a fully disordered school and 1 denoting perfectly ordered school.

From the spatial positions, we also compute the distance to the nerest-neighbour for each fish, and compute the average NND at each time point.

We plot histograms of speed, polarisation and neareast-neighbour distance, and make inferences based on these histograms. Assuming that the time-series of these quantities follow broad stationarity assumptions, such histograms from time series data are a well-accepted estimates of the steady-state distributions of the respective quantities.

#### Group splitting

For each trial of each composition, we compute the proportion of the frames with group splitting (see *Data cleaning and preprocessing* ) as the number of frames with splitting divided by total number of frames. For calculating group properties, we discard the frames with group splitting. Since the distributions of various group properties (e.g. polarization, cohesion, speed) are skewed, we report the mode, median, and standard deviation as appropriate.

#### Spatial sorting

To quantify the degree of assortment of species within the groups, we calculate strong sorting percent which is the measure of how often conspecifics form a highly connected part of the heterospecific group (which we then compare to the same metric computed for a random group, see below). Although the visual inspection reveals minor colour differences between two fish species in our videos (we recall that the videos are taken from the top of the fish tank, see Fig. 1 A-B), the tracking software could not distinguish species via automated tracking process. Therefore, we resorted to manual identification of species for 1000 frames for each heterospecific treatment (200 frames from each replicate, with frames equally spaced in time). For each frame, we find the size of the largest connected subcomponent of conspecifics where two individuals are considered connected if they are Voronoi neighbours. We compute the frequency of frames in which the size of the connected subcomponent of conspecifics is equal to the number of conspecifics in the group. We then calculate the null model for the same data, where positions of individuals from the real frame is randomly shuffled and the sorting is computed for this null frame; this process is repeated 1000 times to obtain the null distribution of sorting values. We calculate how far the strong sorting percent lies from the null distribution. To test this we calculate Zscore which is real mean-null mean/SD of null mean. If the Z score is greater than 2, we consider that the sorting is significantly higher from the null distribution.

### Individual-based model of mixed species schooling

We develop an individual-based spatial model of collective movement that explicitly treats speed as a dynamical variable to account for the observed speed variability in our data. The model is an extension of recent literature [30, 27, 29, 42], with the addition of a new speed-matching interaction, which is key to reproducing the observed results.

#### Basic model

The model consists of N fish, each described by its 2-dimensional position **r**_*i*_(t), speed v_*i*_(t) and its orientation angle φ_*i*_(t). We denote the orientation vector by **e**_*i*_(t) = (cos φ_*i*_(t), sin φ_*i*_(t)) and the velocity vector by **v**_*i*_(t) = v_*i*_(t) **e**_*i*_(t).

In the absence of interactions, every fish has a preferred speed v_0_, to which it relaxes with a relaxation rate *µ*_ind_. Further, its movement (both direction and speed) is affected by noise fluctuations. In addition to this, a fish interacts with its Voronoi neighbors in the following ways:

- *Speed matching*. The fish tries to match its swimming speed to the average velocity of its Voronoi neighborhood, with a rate *µ*_gr_.
- *Alignment or direction matching*. The fish tries to align to the average direction of its neighbourhood, with a rate *µ*_al_.
- *Distance regulation (attraction/repulsion)*. The fish is attracted towards neighbours, and repelled from neighbours that are too close, with an attraction/repulsion strength *µ*_at_.

Alignment, attraction and repulsion interactions are common features of many models of collective movement [11, 12], and some recent models have also incorporated speed variability [30, 27, 29, 42]. A novel feature of our model is the *speed matching* interaction, through which fishes match their swimming speed to match their neighbours. These speed matching interactions are necessary to reproduce the bimodal speed dynamics observed in real data.

The overall dynamics of fish i can be written in terms of the following Langevin equations for v_*i*_(t) and φ_*i*_(t) (in an Ito sense):

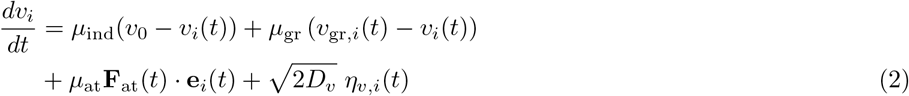

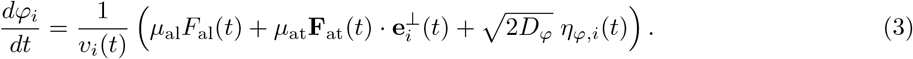

Here, v_gr,*i*_ is the centroid speed of the local neighbourhood of i, defined as

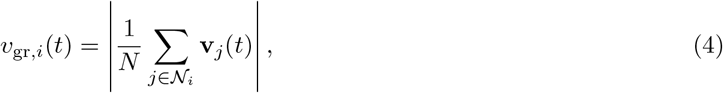

where *N*_*i*_ represents the Voronoi neighbourhood of fish i. Terms F_al_ and **F**_*d*_(t) are the alignment and distance regulation *‘forces’* respectively, given by,

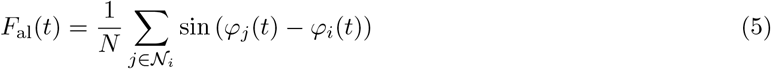

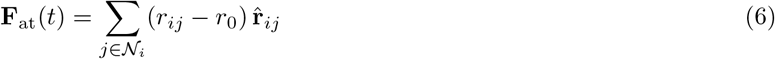

where r_0_ is the preferred distance, **r**_*ij*_ = **r**_*j*_ − **r**_*i*_, r_*ij*_ = |**r**_*ij*_| and 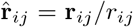. The alignment term F_al_ only acts on the direction φ_*i*_, while the attraction term **F**_*d*_ is resolved along the parallel and angular direction as **F**_*d*_ · **e**_*i*_ and 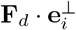 respectively.

The coefficients *µ*_ind_, *µ*_gr_, *µ*_al_ and *µ*_at_ govern the relative strengths of the speed relaxation and various interaction terms respectively. η_*v*_(t) and η_*φ*_(t) are Gaussian white noise terms, and D_*ν*_ and D_*φ*_ are the noise strengths (assumed to be same for both v_*i*_ and φ_*i*_ for simplicity).

#### Mixed species model

For mixed species version of the model, the parameter values for v_0_, r_0_, *µ*_ind_, *µ*_gr_, *µ*_al_, *D*_*v*_ and *D*_*φ*_ can in principle be different between the two species, making the interactions *non-reciprocal* in general [43]. In addition, the distance regulation term *µ*_at_ can be different for inter-versus intra-species interactions. The parameter values used in our simulations are summarized in Table. 1. The values for the preferred speed v_0_ and repulsion zone r_0_ were set based on the modes of the speed and NND distributions respectively. Other parameters—specifically the interaction rates—were tuned to qualitatively reproduce the observed patterns in data.

#### Simulations

We simulated the dynamics of fish schools for the same group size (N = 16) and compositions as in the experiments. We simulated the individual dynamics by simulating Eqs. 2,3 using Euler-Maruyama integration, with a timestep dt = 0.001s. Simulations were done for T = 20,000 time steps each, for a grid of 100 initial conditions, with initial polarization ranging from 0 to 1 and initial speed ranging from 0 to 9.6 BL/s. The simulation results were pooled together, and relevant metrics (viz. polarization histogram, speed histogram, and sorting probabilities) were computed from the pooled data.

## Ethics statement

This research was approved by the Central Animal Facility at the Indian Institute of Science (CAF/Ethics/958/2023).

## Aknowledgements

We thank Kavita Isvaran for inputs on data analysis, Hari Sridhar for general discussions, Thibaut Arnoulx de Pirey for discussions on an early version of the model and Manjunath for assistance with fish lab maintenance. JT was supported by the Prime Minister Research Fellowship, Ministry of Education, Government of India.

## Supplementary Information

### Variation in body length

The variation in body length (BL) of individuals within a trial computed via the coefficient of variation (CV) for each trial. The average CV across all trials was 6%, indicating that the body length of individuals within trials had very low variance compared to mean. The table below captures the mean body length for each trial along with the the CV.

**Table S1.** Mean body length (in cm) for each trial with Coefficient of variation (CV) in bracket.

| Replicate | RB16 | RB12+TB4 | RB8+TB8 | RB4+TB12 | TB16 |
| --- | --- | --- | --- | --- | --- |
| 1 | 3.2 (5%) | 3.5 (6%) | 3.5 (10%) | 3.6 (7%) | 3.8 (5%) |
| 2 | 3.3 (6%) | 3.6 (7%) | 3.6 (9%) | 3.6 (6%) | 3.6 (8%) |
| 3 | 3.5 (6%) | 4.2 (5%) | 3.9 (10%) | 3.5 (5%) | 3.7 (6%) |
| 4 | 4.4 (4%) | 3.9 (5%) | 4.0 (7%) | 3.8 (7%) | 3.6 (3%) |
| 5 | 4.4 (2%) | 4.2 (7%) | 4.1 (7%) | 3.9 (8%) | 3.7 (4%) |

### Group Splitting

**Figure S1.**
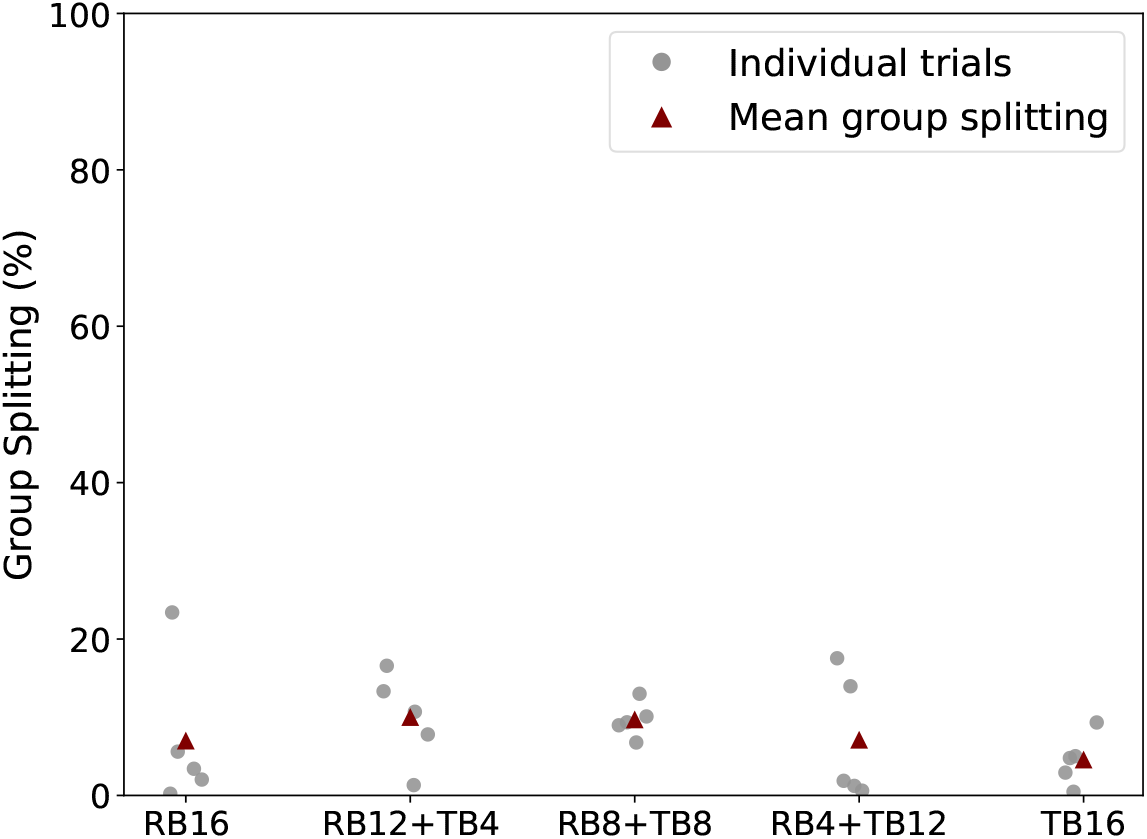
We considered a group as split when even a single individual was more than 6 times average NND far from its nearest neighbour. Each grey dot represent percentage of duration over which the group was split for a single trial whereas the maroon triangle represents the mean computed across replicates for that experimental treatment. We find that mean splitting across all experimnetal conditions (conspecific and heterospecific) was less than 10% of the duration of the experiments, suggesting that groups were cohesive for the major duration of the experiments.

### Lack of trends in speed distributions across replicates and time

**Figure S2.**
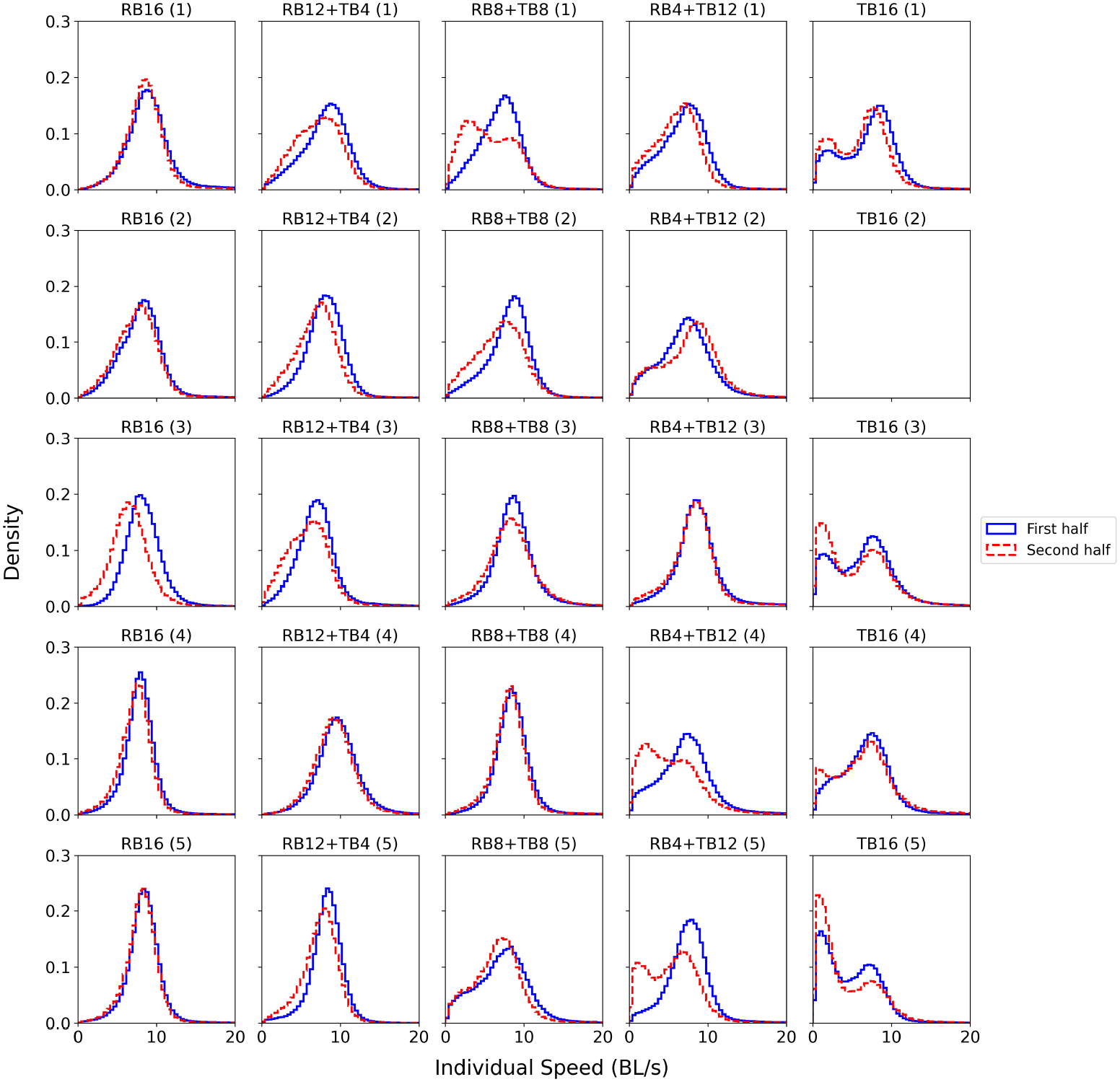
Individual speed distributions and speed matching across replicates. Tiger barbs show bimodality in both first and second half of the trial across all replicates, which indicates that bimodality is not merely a consequence of fish slowing down in the second half of the trials (see last column). The trial to trial variation in bimodality of tiger barbs was also present in model simulations, where sometimes the slow mode is higher and vice verse. Note that the second replicate of TB 16 was discarded due to low quality of tracking. Rosy barbs show unimodal speed distribution across all replicates. Moreover, nearly all replicates show qualitatively similar features across all experimental conditions, the observed results are unlikely to be a consequence of variation in fish behavior across replicates. In mixed species groups the slow swimming mode of tiger barbs is sometimes present in the second half of the trial, but only in three of 15 trials, thus suggesting that variability across trials is low.

### Summary statistics of group properties

**Table S2.** Summary statistics of nearest neighbor distance (in units of BL), individual speed (BL/s), and group polarization across treatments. Each value for the group properties is calculated from the pooled data across all replicates of the respective treatment.

| Treatment | Nearest Neighbor Distance (BL) |  |  |  | Individual Speed (BL/s) |  |  |  | Group Polarization |  |  |  |
| --- | --- | --- | --- | --- | --- | --- | --- | --- | --- | --- | --- | --- |
|  | Mode | Median | Mean | SD | Mode | Median | Mean | SD | Mode | Median | Mean | SD |
| RB16 | 0.53 | 0.55 | 0.58 | 0.20 | 8.10 | 8.00 | 8.00 | 3.00 | 0.98 | 0.97 | 0.94 | 0.09 |
| RB12+TB4 | 0.55 | 0.59 | 0.61 | 0.19 | 8.30 | 8.00 | 8.00 | 3.00 | 0.98 | 0.97 | 0.92 | 0.12 |
| RB8+TB8 | 0.55 | 0.60 | 0.62 | 0.18 | 8.30 | 7.70 | 7.60 | 3.40 | 0.98 | 0.97 | 0.90 | 0.15 |
| RB4+TB12 | 0.55 | 0.56 | 0.57 | 0.14 | 7.90 | 7.40 | 7.40 | 4.00 | 0.98 | 0.97 | 0.87 | 0.20 |
| TB16 | 0.55 | 0.56 | 0.57 | 1.50 | 1.10 & 7.30 | 6.00 | 5.90 | 3.90 | 0.98 | 0.95 | 0.80 | 0.25 |

### Z Score Table for Spatial Sorting

**Table S3.** Z-scores table for Spatial Sorting (See Methods)

| Composition | Null Mean | Real Mean | Z score | P value |
| --- | --- | --- | --- | --- |
| RB12+TB4 (TB) | 17 | 53 | 13.8063 | < 0.0001 |
| RB8+TB8 (TB) | 50 | 78 | 7.6010 | < 0.0001 |
| RB4+TB12 (RB) | 17 | 30 | 4.8345 | < 0.0001 |
| RB8+TB8 (RB) | 50 | 63 | 3.4935 | < 0.001 |

### Speed-Polarization Relationship

**Figure S3.**
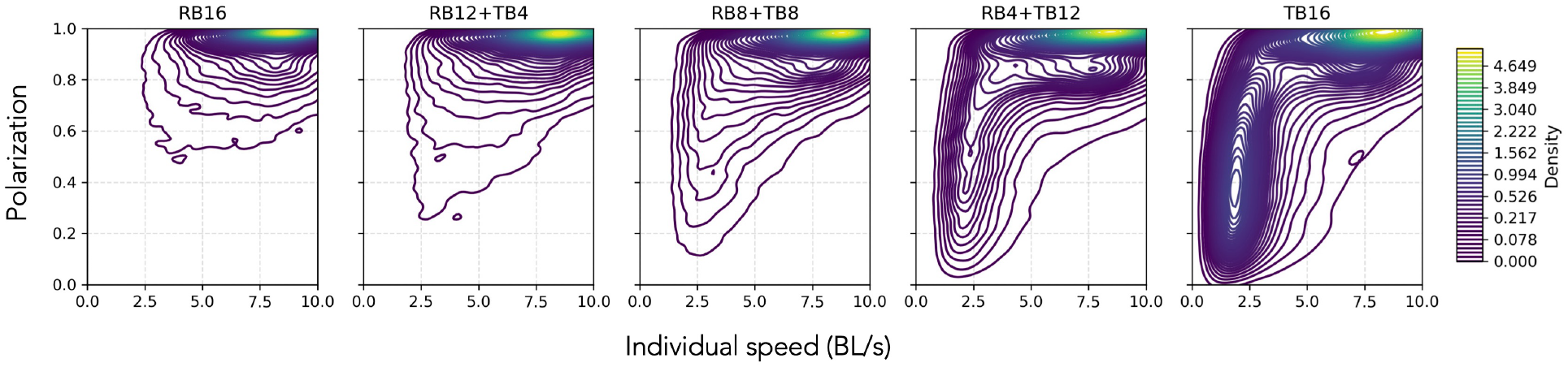
Speed and polarization relationship in the experimental data across mixed species composition. Speed and polarization show a nonlinear relationship in the experimental data, where high speed is associated with high polarization. The bimodality of speed is evident for conspecific groups of tiger barbs (TB 16), where slow speed is associated with low polarisation or a disordered state whereas high speed is associated with high polarisation or an ordered state of fish schools.

### Model shows high polarization across different composition of the two species

**Figure S4.**
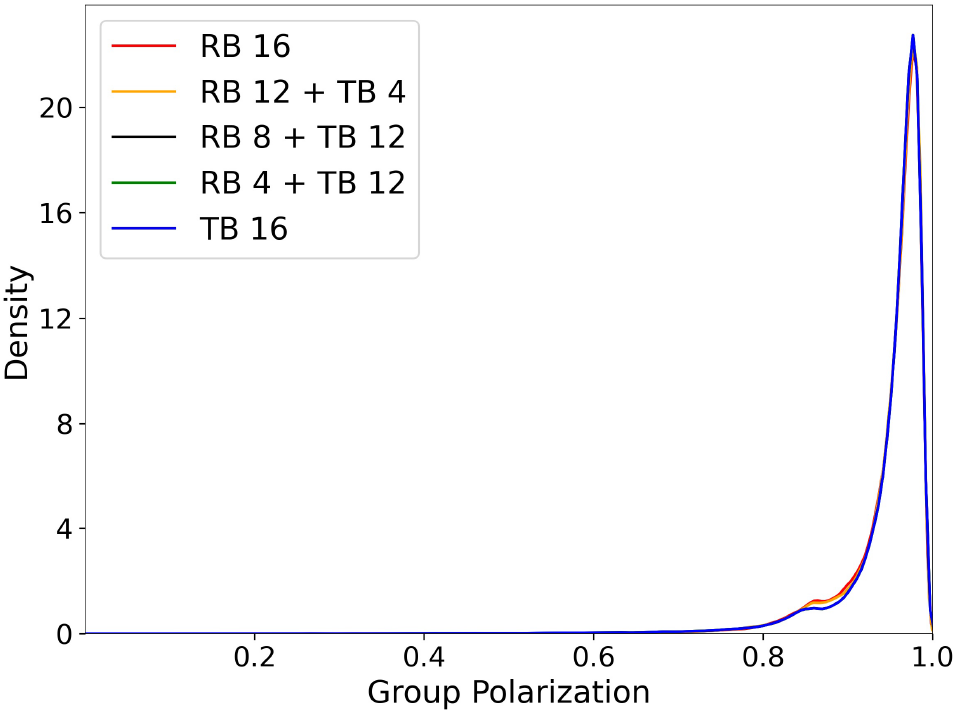
Model shows high polarization across different composition of the two species All the compositions between the two species show high polarization in the model. For other parameters values refer to Table. 1 in Method section.

### Individual speed distributions for different values of strength of speed matching (*µ*_*gr*_) in the model

**Figure S5.**
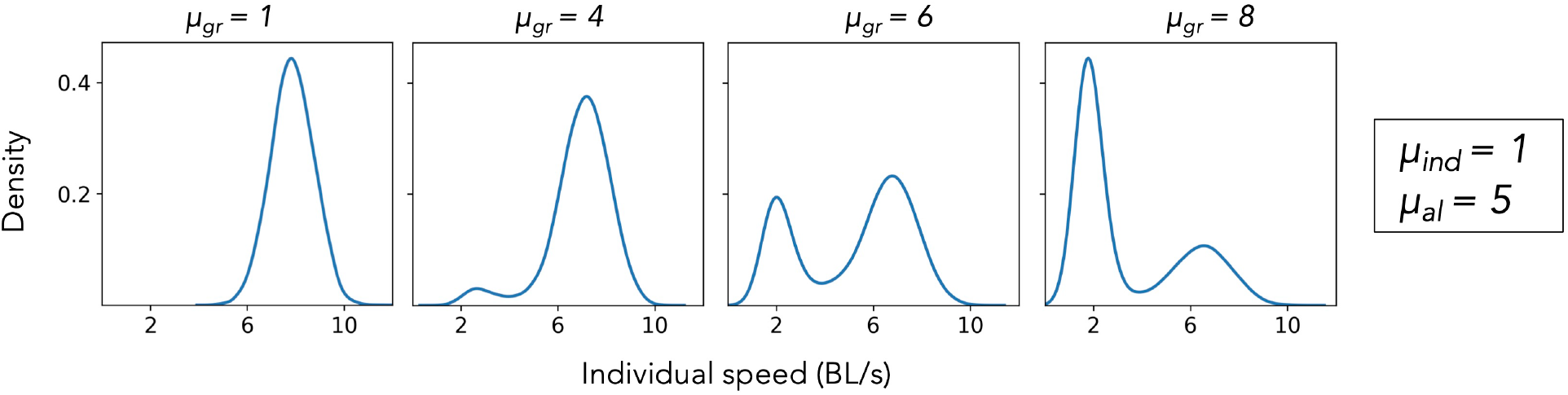
Individual speed distribution across changing strength of speed matching (*µ*_*gr*_ ) For low values of *µ*_*gr*_, we find unimodal distribution of speed mimicking rosy barbs, while for the higher values we find bimodality in speed distribution mimicking tiger barbs, with *µ*_*gr*_ = 6 being the best match for tiger barb distribution. The ratio between strength of relaxation speed (*µ*_*ind*_) and speed matching (*µ*_*gr*_) also defines the distribution of speed in the model. If the two parameter values are comparable, the distribution is unimodal. However, when (*µ*_*gr*_) is much greater than (*µ*_*ind*_), we find bimodal speed distribution. For other parameters values refer to Table. 1 in Method section.

### Spatial sorting across varying intra-specific attraction strength 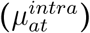 in the model

**Figure S6.**
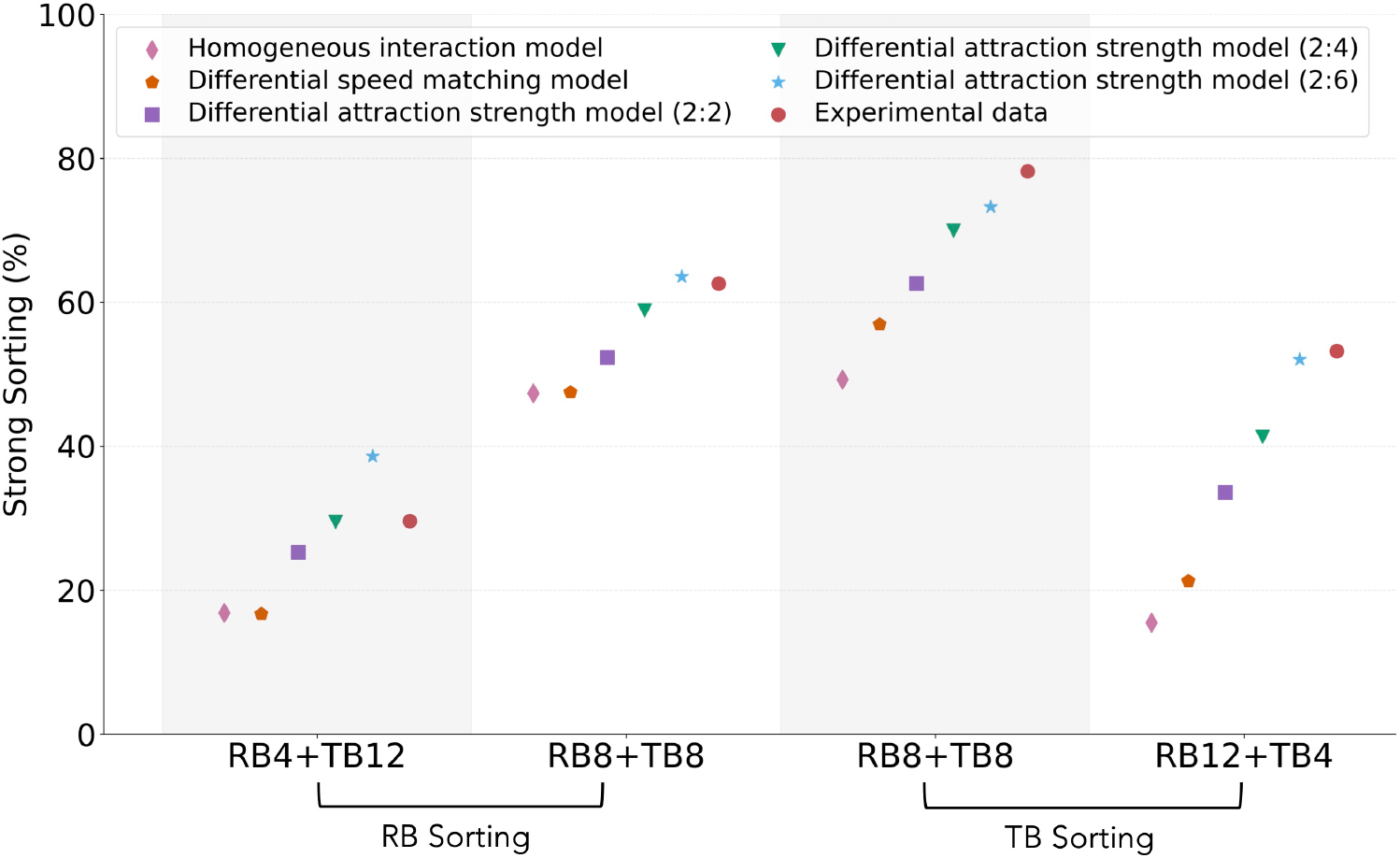
Strong sorting percent across changing intra-specific attraction strength 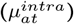.In the homogeneous interaction model, all parameter values are equal for both species, representing a null model for sorting. The Differential-speed-matching model has distinct strengths of speed matching (*µ*_*gr*_ ) for the two species, which explains the speed distribution but not the spatial sorting in the experimental data. Finally, we have the Differential attraction strength model, a modification of the differential speed matching model, where two species can also have different intra-specific attraction strength 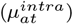. The values in brackets in legend shows the ratio of the attraction strengths 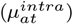 between rosy barbs and tiger barbs. We find that the differential attraction model shows sorting patterns similar to experimental data when 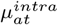 for tiger barbs is three times stronger than that of rosy barbs (2:6). For other parameter values, refer to Table. 1 in the Methods section.

## Notes

### Competing Interest Statement

The authors have declared no competing interest.

